# Comparative Analysis of Test Tube and Volumetric Drinking Monitor Methods in Voluntary Ethanol Consumption in Female Mice for Prenatal Alcohol Exposure

**DOI:** 10.1101/2025.06.10.658718

**Authors:** Arasely M. Rodriguez, Korbin C. Bauer, Lee Anna Cunningham

**Affiliations:** Department of Neurosciences, University of New Mexico Health Sciences Center

**Author notes:** Corresponding Author: Lee Anna Cunningham, Dept. Neurosciences, MSC08 4740, 1 University of New Mexico, Albuquerque, NM 87131-0001. Funding Sources: NIAAA 5R01AA027462, NIAAA 1F31AA031920; NIAAA 2P50AA022534 and NIGMS 5P20GM109089.

## Abstract

**Objective:** Fetal alcohol spectrum disorders affect approximately 1 in 20 school age children in the United States of America. To study fetal alcohol spectrum disorders, mouse models are commonly used. Of the many approaches of gestational exposure, voluntary drinking paradigms represent the most similar mechanism of drinking as human exposure. These exposures can be done through low-tech solutions such as test tubes (TT), or more high-tech methods such as a volumetric drinking monitor (VDM). Here were compare the TT method and the VDM directly, to evaluate their effect on female mouse drinking.

**Method:** We adapted a drinking in the dark, active cycle, limited access (4 hr.) voluntary drinking paradigm first described by Brady et al. (2012) to test tubes and the volumetric drinking monitor. 8 mice were placed in either drinking method and we evaluated their drinking volume and blood alcohol concentrations (BACs). We compared the values for each group using t-tests.

**Results:** After 2 weeks of drinking 10% ethanol with 0.4% saccharine, BACs were not significantly different [t(14)=0.2681, p=0.7935] between the VDM (81.56 ± 21.16 mg/dL) *vs*.TT (73.14 ± 23.20 mg/dL) groups. Calculated intake of ethanol (g/kg) on the day of blood draw for BAC analysis was also not significantly different [t(14)=0.4308, p=0.6732] between VDM (2.985 ± 0.4127) vs.TT (3.260 ± 0.4863; Fig 1B) groups.

**Conclusions:** Test tube or VDM resulted in similar average daily ethanol consumption and resultant BACs in female mice

## Introduction

Approximately 1 in 20 school children in the United States has a fetal alcohol spectrum disorder (FASD) (May et al., 2018). Gestational alcohol exposure can lead to a broad spectrum of neurodevelopmental disabilities collectively defined as fetal alcohol spectrum disorders (FASDs). Gestational alcohol exposure was first associated with neurodevelopmental disabilities in 1968 by Lemoine et al. Later, Jones & Smith (1973) characterized fetal alcohol syndrome (FAS) as the most severe sub-group of FASDs, which results from high doses of alcohol during gestation, and includes facial dysmorphology, growth restriction and disorders of the central nervous system. Since then, research has shown that exposure to more moderate doses of alcohol during gestation can result in a spectrum of FASDs that encompass a range of deleterious effects on brain development without facial dysmorphologies, associated with long-term behavioral and cognitive impairments (Caputo et al., 2016; Hellemans et al., 2010; Mattson et al., 2019; Popova et al., 2021). Average life expectancy of the most severe form of FASD, FAS, is 34 years of age (Thanh & Jonsson, 2016).

**Fig 1.**
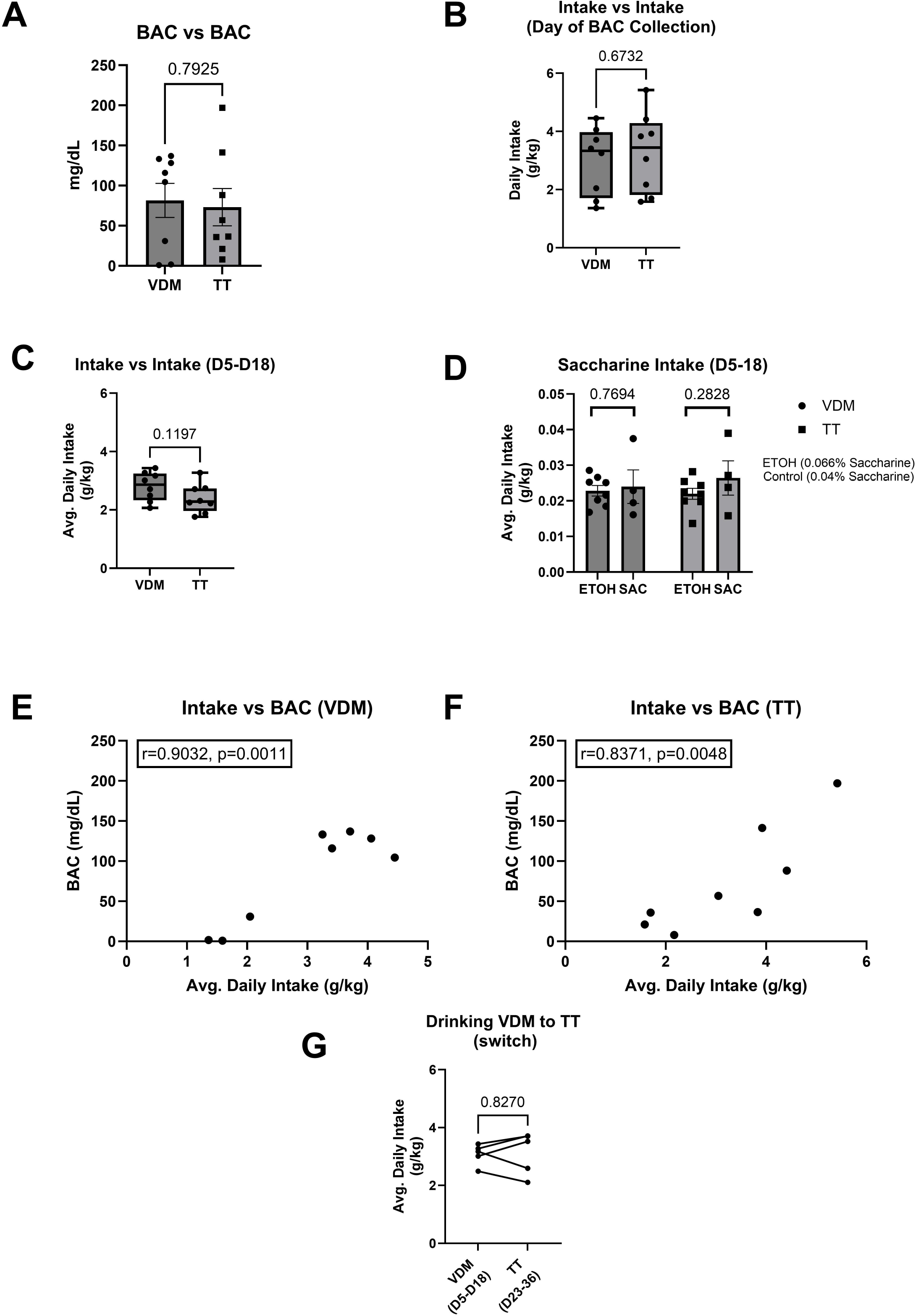

A wide range of pre-clinical animal models of FASD have been established to understand the neurodevelopmental effects of gestational alcohol. These experimental models utilize species ranging from the microscopic, C. elegans, to mammals and non-human primates. However, mouse models represent the most ubiquitous species for modeling prenatal alcohol exposure (PAE) (Almeida et al., 2020). Exposure models also vary widely, including variations in timing during pregnancy and modes of alcohol administration, which can include alcohol administration by intragastric gavage (IG), vapor inhalation, intraperitoneal injection (IP), or voluntary oral consumption. Each of these administration methods have their advantages and limitations in mimicking human alcohol consumption during pregnancy. Methods such as IG and IP offer more controlled delivery of alcohol and higher blood alcohol concentrations (BACs), but add an additional component of stress (Almeida et al., 2020; Patten et al., 2014). Voluntary oral consumption mimics human consumption and is less stressful, but also results in lower BACs that model more moderate levels of alcohol exposure (Almeida et al., 2020; Patten et al., 2014).

A commonly used model of voluntary alcohol consumption during pregnancy, originally described by Brady et al. (2012), is a voluntary limited access drinking-in-the-dark paradigm. In this paradigm, alcohol is provided in the drinking water for a limited period (4 hours per day) during the animals awake (dark) cycle using a system of test-tubes with a ball bend sipper, with amounts of ingested alcohol calculated volumetrically at the end of each drinking period.

Although the BACs achieved by this model are low (Licheri et al., 2023) to moderate (Brady et al., 2012), this paradigm results in long-lasting neurological impairments in offspring that mimic many of those described in FASD (Gustus et al., 2020; Valenzuela et al., 2012). More technological precise methods of oral drinking during PAE exposures have been implemented through the use of the automated volumetric drinking monitor system (VDM; Myrick et al., 2024). The VDM is an automated approach that utilizes a micropump to dispense precise amounts of liquid. While the VDM approach has many advantages for alcohol studies, including the ability to continuously and precisely monitor rate of intake throughout the entire exposure period, it is also an expensive apparatus that may not be necessary for most alcohol studies. The present report describes a side-by-side comparison of the test tube vs. VDM alcohol administration methods to determine whether there are any differences regarding levels of alcohol consumed or BACs achieved in female mice of reproductive age.

## Methods

### Animals

Because our laboratory utilizes transgenic Nestin-CreER^T2^:tdTomato reporter mice to investigate the effects of developmental alcohol on adult hippocampal neurogenesis, we utilized this strain in the current study. This bitrangenic strain is maintained on a C57BL/6J background (Lagace et al., 2007; Madisen et al., 2010). These mice were taken from breeding colonies maintained by the University of New Mexico Health Sciences Center Animal Resource Facility. Mice were housed under reverse 12:12 hour light cycle with lights off at 0800 in a humidity and temperature-controlled room. Unless stated otherwise mice had *ad libitum* access to water and standard animal chow (Teklad 2920X). Tamoxifen was not administered as this study focused on alcohol consumption. Animal procedures were approved by the University of New Mexico Institutional Animal Care and Use Committee in accordance with NIH policies on care and use of laboratory animals. For this study only female mice of breeding age were used, given our interest in gestational exposure paradigms.

### Limited Access Drinking-in-the-Dark Prenatal Alcohol Exposure

The paradigm described herein was adapted from Brady et al. (2012). Female mice of reproductive age, (average 152 days old), were maintained on a reverse light dark cycle. Mice were single-housed and acclimated to reverse light for one week. After acclimatization, water bottles were replaced with bottles containing 5% ethanol in 0.066% (w/v) saccharine or 0.04% saccharine in water alone (control). Although saccharine is necessary for ethanol consumption in the NestinCreERT2:tdTom strain; we reduced the concentration of saccharine in control solutions to ensure no differences in liquid consumed. Mice were initially provided 5% ethanol/saccharine or saccharin alone for 4 consecutive days to establish drinking, followed by 10% ethanol/saccharine or saccharine alone throughout the next 25 days. Water was provided to all mice outside of the 4 hour drinking period.

### Test Tube (TT) Sippers

55 mL, 25 × 150 mm test tubes which have a size 4 rubber stopper affixed with a bend ball sipper (Anacare, BW-201F) were used. Tubes were filled 1 inch below the rim with solutions of ethanol/saccharine or saccharine-only as described above. Tubes were weighed before and after the drinking period to calculate volume reduction following the drinking period. Two control TTs filled with each solution were placed in empty cages and removed after the drinking period ended; these were then weighed and used to account for any leakage. After the 4 hour drinking period, all TTs were replaced with standard water bottles.

### Volumetric Drinking Monitor (VDM)

The VDM was purchased from Columbus instruments (Columbus, OH). This system dispenses 20 microliters of fluid for each mouse lick. The volume of liquid dispensed was digitally recorded by VDM software on a continuous basis and stored as consecutive 10-second bins over the entire 4 hour exposure period (1440 bins), as previously described by (Myrick et al., 2024). The liquid dispensed by the VDM is drawn from reservoirs of either ethanol-saccharine or saccharine-only solutions as described above. The system was calibrated per manufacturer instructions.

### Blood Alcohol Concentration (BAC)

Blood samples were drawn from the peri-orbital sinus from the alcohol-exposed mice after 14 days of drinking, immediately following the 4 hour exposure period (Boehm et al., 2008). Plasma was isolated and kept frozen at −20C until analyzed. Serum alcohol levels were measured using an Analox Alcohol Analyzer (Analox Instruments, Lunenburg, MA) following manufacturer instructions, as previously described in Lee et al. (2000)

### Switching Mice from VDM to TT

A small cohort of mice were switched from the VDM group to the TT method on day 22 of drinking after the 4 hour drinking period on that day. After 7 days of acclimation to the new drinking apparatus, we monitored ethanol intake on the TT relative to the prior VDM intake.

### Statistical Analysis

All data were subjected to statistical analysis using Graphpad Prism (10.4.1) software. Data are described as means ± SD. Two-tailed t-tests were used for comparison with p-values<0.05 considered significant. Error bars in figures represent SEM.

## Results

To compare the traditional test tube method with the modern volumetric tracking system, 8 female mice were assigned to test tube (TT) or volumetric drinking monitor (VDM) groups. All mice were acclimated for the same period of time to the reverse light dark cycle and provided with the same daily 4 hour exposure (4 days at 5% ETOH and 25 days at EtOH 10%). After 14 days of drinking, blood was collected from each group and BACs (mg/dL) were determined. As shown in Fig. 1A, BACs were not different between the VDM (81.56 ± 21.16 mg/dL) *vs*.TT (73.14 ± 23.20 mg/dL) groups [t(14)=0.2681, p=0.7935]. To determine if there was a difference in ethanol intake, we also calculated the intake of ethanol as g/kg. On the day of BAC collection, we found no significant difference between VDM (2.985 ± 0.4127 g/kg) vs.TT (3.260 ± 0.4863 g/kg) groups [t(14)=0.4308, p=0.6732] (Fig. 1B). The average daily intake of ethanol (g/kg) over 14 consecutive days of drinking 10% ethanol, (i.e., day 5 through day 18) was not significantly different [t(14)=1.657, p=0.1197] between VDM (2.803 ± 0.1751 g/kg) and TT (2.393 ± 0.1748 g/kg; Fig. 1C).

To ensure that total saccharine consumption also did not vary between control and ethanol groups, daily saccharine intake was calculated as g saccharine consumed/kg body weight. The saccharine intake in VDM mice was not significantly different [t(10)-1.135; p = 0.7694] between the saccharine control (0.02398 ± 0.009421 g/kg) and the ethanol (0.02283 ± 0.004170 g/kg; Fig. 1D) groups. Similarly, the saccharine intake in the TT mice was not significantly different [t(10)=1.135, p=0.2828] between the saccharine controls (0.02644 ± 0.009621 g/kg) and the ethanol mice (0.02197 ± 0.004375 g/kg; Fig. 1D).

Ethanol intakes were compared to BACs to determine whether they were closely correlated. There was a significant positive correlation between BAC and ethanol intake both in the VDM mice [r(6)=0.9032, p=0.0011; Fig. 1E] and TT mice [r(6)=0.8157, p=0.0048; Fig. 1F].

On day 23 of drinking, 4 high drinking mice in the VDM group were switched to test tubes to see if this affected intake. The average daily intake of ethanol was not significantly different [t(4)=0.2333, p=0.8270] between their time drinking on the VDM (3.074 ± 0.3604 g/kg) vs. the TT (3.126 ± 0.7323 g/kg; Fig. 1G)

## Discussion

The VDM was first implemented in drinking studies in 1987, by Blanchard et al. where it was used to evaluate the effect of ethanol on social structure in laboratory rats. With the VDM, Blanchard et al. were able to describe frequency of ethanol consumption based on the pattern of licks. More contemporary studies have used the VDM to evaluate precise liquid volumes to describe more nuanced drinking behavior, such as front loading in the binge drinking models, where animals rapidly consume ethanol early in the exposure period (Maphis et al., 2022).

Here we performed a side-by-side comparison to demonstrate that the VDM and TT produce similar BACs in female mice. Likewise, the ethanol intake on the day BACs were collected was not different between the VDM and the TT. The average daily ethanol intake across two weeks was also not different between VDM and TT. These findings indicate that VDM and TT achieve similar exposure levels in mice. Furthermore, the relationship between the ethanol intake and the BACs were strongly correlated, with the VDM being slightly more tightly correlated compared with the TT. Nevertheless, the high correlation suggests that the total ethanol intake could be used as a substitute for BACs in both methods. Additionally, we showed that saccharine intake between the ethanol and control mice (control) were similar in both methods. This is important aspect to control for since many strains of mice, including those used in the present study, will not drink ethanol in water alone. Overall, these data demonstrate comparable daily ethanol intake is achieved using either the VDM or the TT administration method.

Although the VDM is a more technically accurate system for continuous monitoring, it has both its advantages and disadvantages for routine alcohol studies. One advantage is that the VDM monitors drinking volumes continuously for a defined period of time which can be used to evaluate the pattern of drinking of the mice. Importantly, the pattern of alcohol consumption may influence neurobehavioral outcomes in offspring (Myrick et al., 2024). However, the VDM is also relatively expensive and requires a greater investment of logistical factors, including ongoing calibrations and the need for specialized components. While the TT method is less precise, it is readily accessible and cost effective. Although there are differences in data resolution monitoring comparing TT to VDM, here we show that the average daily alcohol consumption and resultant BACs are similar regardless of TT vs. VDM administration in this limited access drinking-in-the dark paradigm. Furthermore, the two systems are interchangeable in this regard, since switching mice from VDM to TT during the exposure period did not alter daily ethanol consumption. To summarize, for routine alcohol exposure paradigms, there is no advantage of VDM over TT as both provide similar intake and BACs.

## Acknowledgements

We thank Dr. Elif Tunc-Ozcan for helpful feedback and insightful comments on drafting this manuscript. We also thank the New Mexico Alcohol Research Center staff for their technical assistance. This research was supported by NIAAA 5R01AA027462, NIAAA1F31AA031920; NIAAA 2P50AA022534 and NIGMS 5P20GM109089.

## References

Almeida, L., Andreu-Fernández, V., Navarro-Tapia, E., Aras-López, R., Serra-Delgado, M., Martínez, L., García-Algar, O., & Gómez-Roig, M. D. (2020). Murine Models for the Study of Fetal Alcohol Spectrum Disorders: An Overview. Frontiers in Pediatrics, 8. 10.3389/fped.2020.00359

Blanchard, R. J., Hori, K., Tom, P., & Blanchard, D. C. (1987). Social structure and ethanol consumption in the laboratory rat. Pharmacology Biochemistry and Behavior, 28(4), 437–442. 10.1016/0091-3057(87)90502-8

Boehm, S. L., Goldfarb, K. J., Serio, K. M., Moore, E. M., & Linsenbardt, D. N. (2008). Does context influence the duration of locomotor sensitization to ethanol in female DBA/2J mice? Psychopharmacology, 197(2), 191–201. 10.1007/s00213-007-1022-6

Brady, M. L., Allan, A. M., & Caldwell, K. K. (2012). A Limited Access Mouse Model of Prenatal Alcohol Exposure that Produces Long-Lasting Deficits in Hippocampal-Dependent Learning and Memory: A LIMITED ACCESS MOUSE MODEL OF PAE. Alcoholism: Clinical and Experimental Research, 36(3), 457–466. 10.1111/j.1530-0277.2011.01644.x

Caputo, C., Wood, E., & Jabbour, L. (2016). Impact of fetal alcohol exposure on body systems: A systematic review. Birth Defects Research. Part C, Embryo Today: Reviews, 108(2), 174–180. 10.1002/bdrc.21129

Gustus, K., Li, L., Newville, J., & Cunningham, L. A. (2020). Functional and Structural Correlates of Impaired Enrichment-Mediated Adult Hippocampal Neurogenesis in a Mouse Model of Prenatal Alcohol Exposure. Brain Plasticity, 6(1), 67–82. 10.3233/BPL-200112

Hellemans, K. G. C., Sliwowska, J. H., Verma, P., & Weinberg, J. (2010). Prenatal alcohol exposure: Fetal programming and later life vulnerability to stress, depression and anxiety disorders. Neuroscience & Biobehavioral Reviews, 34(6), 791–807. 10.1016/j.neubiorev.2009.06.004

Jones, K. L., & Smith, D. W. (1973). Recognition of the fetal alcohol syndrome in early infancy. Lancet (London, England), 302(7836), 999–1001. 10.1016/s0140-6736(73)91092-1

Lagace, D. C., Whitman, M. C., Noonan, M. A., Ables, J. L., DeCarolis, N. A., Arguello, A. A., Donovan, M. H., Fischer, S. J., Farnbauch, L. A., Beech, R. D., DiLeone, R. J., Greer, C. A., Mandyam, C. D., & Eisch, A. J. (2007). Dynamic contribution of nestin-expressing stem cells to adult neurogenesis. The Journal of Neuroscience: The Official Journal of the Society for Neuroscience, 27(46), 12623–12629. 10.1523/JNEUROSCI.3812-07.2007

Lee, S., Schmidt, D., Tilders, F., Cole, M., Smith, A., & Rivier, C. (2000). Prolonged Exposure to Intermittent Alcohol Vapors Blunts Hypothalamic Responsiveness to Immune and Non-Immune Signals. Alcoholism: Clinical and Experimental Research, 24(1), 110–122. 10.1111/j.1530-0277.2000.tb04560.x

Lemoine, P., Harouseau, H., Borteryu, J., & Menuet, J. J. (1968). Les enfants des parents alcooliques. Anomalies observées à propos de 127 cas. https://api.semanticscholar.org/CorpusID:148717406

Licheri, V., Jacquez, B. J., Castillo, V. K., Sainz, D. B., Valenzuela, C. F., & Brigman, J. L. (2023). Long-term effects of low prenatal alcohol exposure on GABAergic interneurons of the murine posterior parietal cortex. Alcohol, Clinical & Experimental Research, 47(12), 2248–2261. 10.1111/acer.15210

Madisen, L., Zwingman, T. A., Sunkin, S. M., Oh, S. W., Zariwala, H. A., Gu, H., Ng, L. L., Palmiter, R. D., Hawrylycz, M. J., Jones, A. R., Lein, E. S., & Zeng, H. (2010). A robust and high-throughput Cre reporting and characterization system for the whole mouse brain. Nature Neuroscience, 13(1), 133–140. 10.1038/nn.2467

Maphis, N. M., Huffman, R. T., & Linsenbardt, D. N. (2022). The development, but not expression, of alcohol front-loading in C57BL/6J mice maintained on LabDiet 5001 is abolished by maintenance on Teklad 2920x rodent diet. Alcoholism, Clinical and Experimental Research, 46(7), 1321–1330. 10.1111/acer.14876

Mattson, S. N., Bernes, G. A., & Doyle, L. R. (2019). Fetal Alcohol Spectrum Disorders: A Review of the Neurobehavioral Deficits Associated With Prenatal Alcohol Exposure. Alcoholism, Clinical and Experimental Research, 43(6), 1046–1062. 10.1111/acer.14040

May, P. A., Chambers, C. D., Kalberg, W. O., Zellner, J., Feldman, H., Buckley, D., Kopald, D., Hasken, J. M., Xu, R., Honerkamp-Smith, G., Taras, H., Manning, M. A., Robinson, L. K., Adam, M. P., Abdul-Rahman, O., Vaux, K., Jewett, T., Elliott, A. J., Kable, J. A., … Hoyme, H. E. (2018). Prevalence of Fetal Alcohol Spectrum Disorders in 4 US Communities. JAMA, 319(5), 474–482. 10.1001/jama.2017.21896

Myrick, A., Jimenez, D., Jacquez, B., Sun, M. S., Noor, S., Milligan, E. D., Valenzuela, C. F., & Linsenbardt, D. N. (2024). Maternal alcohol drinking patterns predict offspring neurobehavioral outcomes. Neuropharmacology, 257, 110044. 10.1016/j.neuropharm.2024.110044

Patten, A. R., Fontaine, C. J., & Christie, B. R. (2014). A Comparison of the Different Animal Models of Fetal Alcohol Spectrum Disorders and Their Use in Studying Complex Behaviors. Frontiers in Pediatrics, 2. 10.3389/fped.2014.00093

Popova, S., Dozet, D., Shield, K., Rehm, J., & Burd, L. (2021). Alcohol’s Impact on the Fetus. Nutrients, 13(10), 3452. 10.3390/nu13103452

Thanh, N. X., & Jonsson, E. (2016). Life Expectancy of People with Fetal Alcohol Syndrome. Journal of Population Therapeutics and Clinical Pharmacology = Journal De La Therapeutique Des Populations Et De La Pharmacologie Clinique, 23(1), e53–59.

Valenzuela, C. F., Morton, R. A., Diaz, M. R., & Topper, L. (2012). Does moderate drinking harm the fetal brain? Insights from animal models. Trends in Neurosciences, 35(5), 284– 292. 10.1016/j.tins.2012.01.006

